# A theory for ecological survey methods to map individual distributions

**DOI:** 10.1101/137158

**Authors:** Nao Takashina, Maria Beger, Buntarou Kusumoto, Suren Rathnayake, Hugh P. Possingham

## Abstract

Spatially-explicit approaches are widely recommended for ecosystem management. The quality of the data, such as presence/absence or habitat maps, affects the management actions recommended, and is, therefore, key to management success. However, available data are often biased and incomplete. Previous studies have advanced ways to resolve data bias and missing data, but questions remain about how we design ecological surveys to develop a dataset through field surveys. Ecological surveys may have multiple spatial scales, including the spatial extent of the target ecosystem (observation window), the resolution for mapping individual distributions (mapping unit), and the survey area within each mapping unit (sampling unit). We developed an ecological survey method for mapping individual distributions by applying spatially-explicit stochastic models. We used spatial point processes to describe individual spatial placements using either random or clustering processes. We then designed ecological surveys with different spatial scales and individual detectability. We found that the choice of mapping unit affected the presence mapped fraction, and the fraction of the total individuals covered by the presence mapped patches. Tradeoffs were found between these quantities and the map resolution, associated with equivalent asymptotic behaviors for both metrics at sufficiently small and large mapping unit scales. Our approach enabled consideration of the effect of multiple spatial scales in surveys, and estimation of the survey outcomes such as the presence mapped fraction and the number of individuals situated in the presence detected units. The developed theory may facilitate management decision-making and inform the design of monitoring and data gathering.

## 1 Introduction

Understanding the spatial characteristics of ecosystems is one of the central challenges in ecology [1]. Such knowledge forms a prerequisite for effective ecosystem management due to an increasing need for spatially explicit approaches in fisheries and wildlife management [2-4] and for the establishment of terrestrial and marine protected areas [5-7].

In ecosystem management, the quality of the data involved in the management decisionmaking, such as presence/absence or habitat maps, affect the management actions recommended [8-10]. Therefore, creating an ecologically and statistically adequate dataset is key to management success. However, available data is often biased and incomplete [8,9], due to, for example, different accessibility to sites [8], existence of the favored study sites [8], and imperfect detectability of individuals [11,12]. These biases hinder the effective implementation of management actions, and may lead to perverse outcomes or wasted management resources. Hence it is important to discuss and benchmark the quality of the spatially explicit data that underlies management decisions.

There is a body of literatures to tackle the challenges of data gathering, including sampling designs for effectively allocating the survey effort under the time and budgetary constraints [13-15], methods for reducing the bias of occurrence data by estimating the detectability of species [12,16-18], and mathematical theory for ecological sampling [19,20]. Although these researches have significantly advanced our insight into ecosystem monitoring and ecological survey, there still remains a question about how we actually design the entire ecological survey to systematically develop dataset through a field survey, as the spatial scale issue, such as how to chose the resolution of a map, is often omitted. This is perhaps because many existing studies consider the space to be sampled implicitly. Presence/absence or habitat map is widely used in ecosystem management [16], where at least three different spatial scales may exist; the spatial extension of the ecosystem under concern, resolution to map the individual distributions, and minimum size of survey units. To systematically gather the spatial data, manager should explicitly take into consideration these three spatial scales, because the manner of the sampling and management outcomes depend on the resolution of a map. For example, in fisheries management, finely implemented fishing quota allocations may result in better management outcomes [7, 21], and this can be done with the distribution map with a high degree of resolution. However, surveying an area at a fine spatial scale is often impractically expensive in the large scale survey, and the choice of resolution itself faces a budgetary constraint. Hence, quantitatively estimating the performance of a sampling method in advance facilitates survey decision making.

In this paper, we develop a theory of ecological survey method for systematically mapping individual distributions by making use of the spatial point processes (SPPs), a spatially explicit stochastic model. The SPPs is widely applied in the study of plant communities [22-25], coral communities [26], and avian habitat selection [27]. Therefore, these are potential target species for the method developed here. However, the method may be suitable for any organism or the location used by an organism (e.g., nesting site) that is relatively sedentary during the time and spatial scale of the field survey so that its spatial distributions can be described by SPPs [28]. In this study, the SPPs describes individual spatial locations by two different processes accounting random or clustering patterns. An ecological survey is then introduced with a set of spatial scales and detectability of individuals. Our spatially-explicit approach is capable of revealing a series of questions important for ecological survey, such as effect of the choice of the spatial scales and spatial distribution patterns of individuals on accuracy of the distribution map. This knowledge enables one to determine the design of an ecological survey beforehand given accuracy of a map required. The developed theory may significantly facilitate management decision making and give solid bases of data gathering.

## 2 Methods

### 2.1 Models of spatial distribution of individuals

To develop a theory of ecological survey to map individual distributions, we explicitly model the spatial distribution patterns of individuals. Spatial point processes (SPPs) [22, 25] provide models to describe such patterns with high flexibility and analytical tractability [24]. Here, we apply the homogeneous Poisson process and the Thomas process, a family of the Neyman-Scott process (Fig. 1).

**Figure 1:**
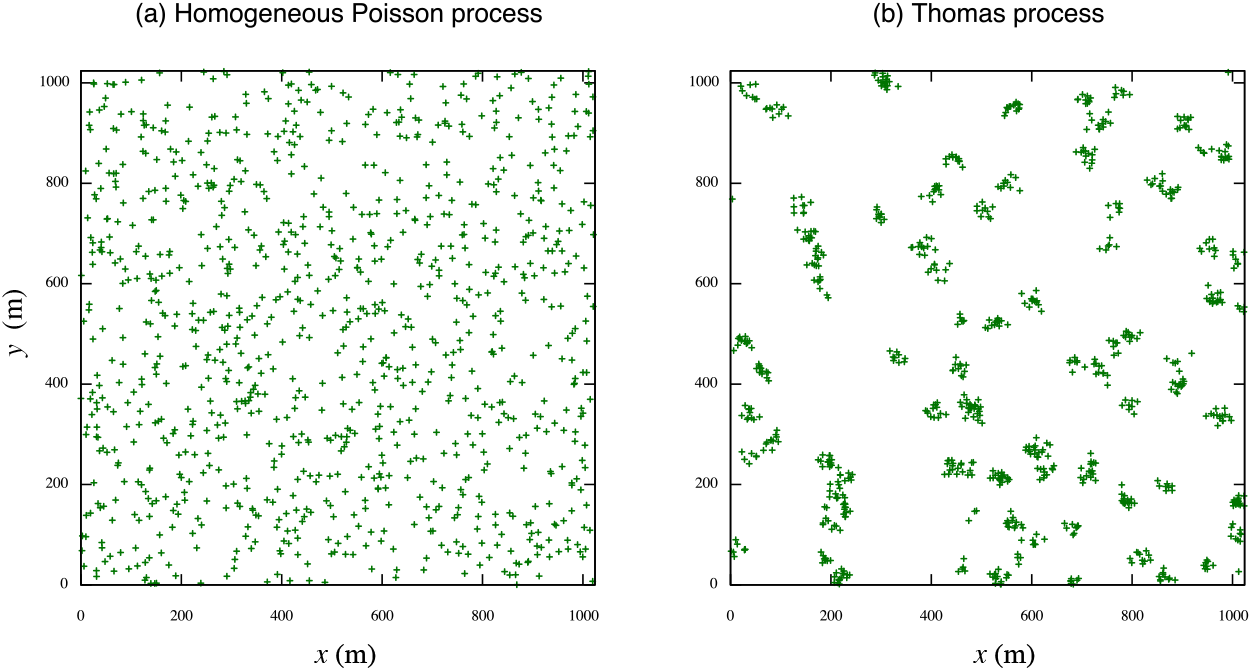
Example of point patterns within a observation window 1024m × 1024m. (a) Homogeneous point process with the intensity ⋋ = 10^−3^; (b) Thomas process with the same intensity value as the homogeneous Poisson process 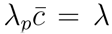, where ⋋_*p*_ = 10^−4^ and 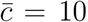. The variance of the bivariate normal distribution σ^2^ = 100. See the text for the interpretations of the parameters.

One of the simplest SPPs is the homogeneous Poisson process where the points (i.e. individuals) are randomly distributed and the number of points of a given region *A, N(A)*, is according to the Poisson distribution with an average *μ_A_:*

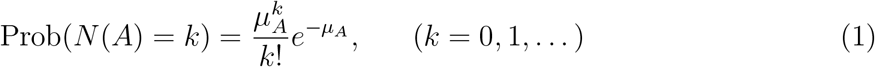

where, *μ_A_* is also regarded as the intensity measure [22,25] described as

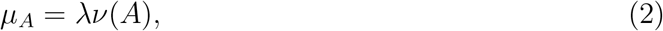

where, *⋋*= *(total points)/(area of concerned region A)* is the intensity in the given region, and *ν*(A) is the area of A.

The Neyman-Scott process [22, 25] provides us more general framework to analyze spatial ecological data and characterize the clustering pattern of individuals [22-25]. By the following three steps, the Neyman-Scott process is obtained:

- Parents are randomly placed according to the homogeneous Poisson process with an intensity *⋋_p_.*
- Each parent produces a random discrete number *c* of daughters, realized independently and identically for each parent.
- Daughters are scattered around their parents independently with an identical spatial probability density, *f* (y), and all the parents are removed in the realized point pattern.

The intensity of the Neyman-Scott process is [25]

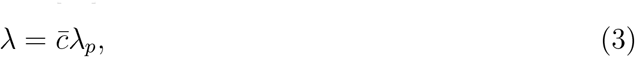

where, 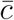 is the average number of daughters per parent. The probability generating functional (pgfl) of the number of daughters within a given region of the Neyman-Scott process is [22,25]

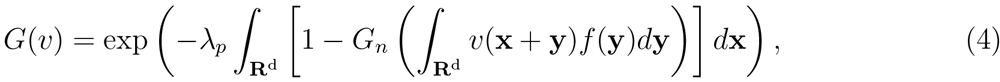

where, 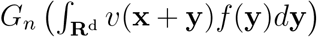 is the probability generating function (pgf) of the random number *c*, the number of daughters per parent.

The Thomas process is a special case of the Neyman-Scott process, where *f*(y) is an isotropic bivariate Gaussian distribution with the variance σ^2^ [25]. We also assume that the number of daughters per parent follows the Poisson distribution with the average number, c. The pgfl of the Thomas process, Eq. (4), within a given region *A* is obtained by substituting the pgf of the number of daughters per parent *G_n_* in Eq. (4). It is obtained, by the given assumptions, as

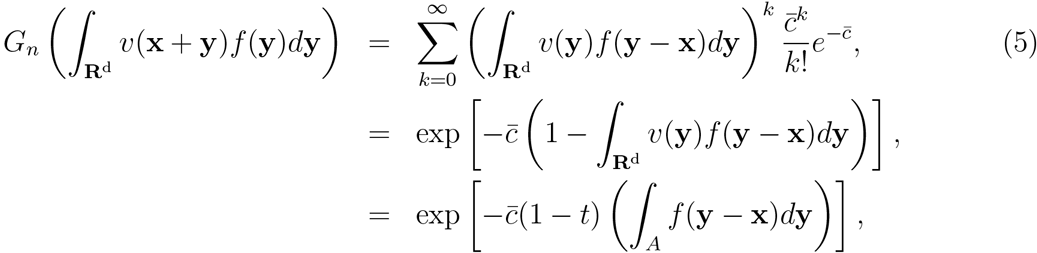

where, to obtain the last line, *v*(y) = 1 − (1 − t)1_A_(y) is used, and here 1_A_(y) is the indicator function. Therefore, the pgfl of the number of daughters within the region *A* of the Thomas process is

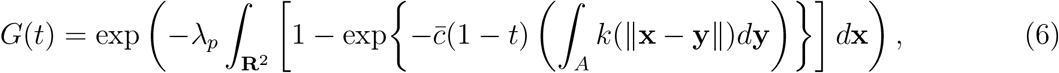

where, *k*(||X − C||) is an isotropic bivariate Gaussian distribution with variance σ^2^,

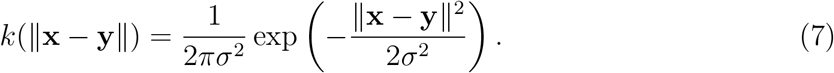

In order to reasonably compare the results of Thomas process with those of the homogeneous Poisson process, we chose the intensity of the Thomas process so as to have, on average, the same number of individuals within the concerned region. Namely, the parameters *⋋_p_* and 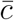 satisfy

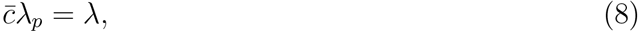

where, the left hand side is the intensity of the Thomas process and the right hand side is the intensity of the homogeneous Poisson.

### 2.2 Design of ecological survey

#### 2.2.1 Survey rules and basic properties

Let us consider the situation where an ecological survey takes place for the purpose of creating a presence/absence map of a given region. A presence/absence map is characterized by the three spatial scales: the *observation window (W*), the spatial scale of ecological survey conducted, the spatial scale of the *mapping unit (M*) defining the resolution of the map, and the spatial scale of the *sampling unit* (S) determining the sampling density within each mapping unit (Fig. 2). We assume the following three key sampling rules.

**Figure 2:**
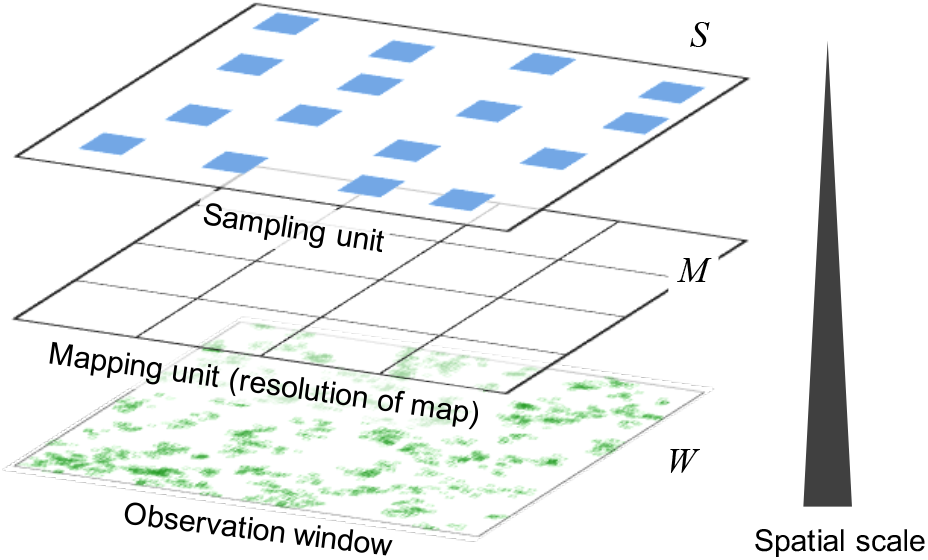
Multiple spatial scales in ecological survey. Each scale is arbitrary determined by managers.

- The observation window, *W*, resolution of map (i.e., scale of the mapping unit), *M*, and sampling unit, *S*, are arbitrary determined, but a single resolution is allowed for each map.
- Every mapping unit is assessed by sampling unit, and sampling location is determined randomly within the mapping unit.
- A mapping unit is recorded as presence if at least one individual is detected in the sampling unit regardless of the number of miss detections. If there is no individual or all individuals are miss detected in the sampling unit, the mapping unit is recorded as absence.

Through the second and third assumptions, changing the scale of the mapping unit affects the obtained presence/absence map (Fig. 3).

**Figure 3:**
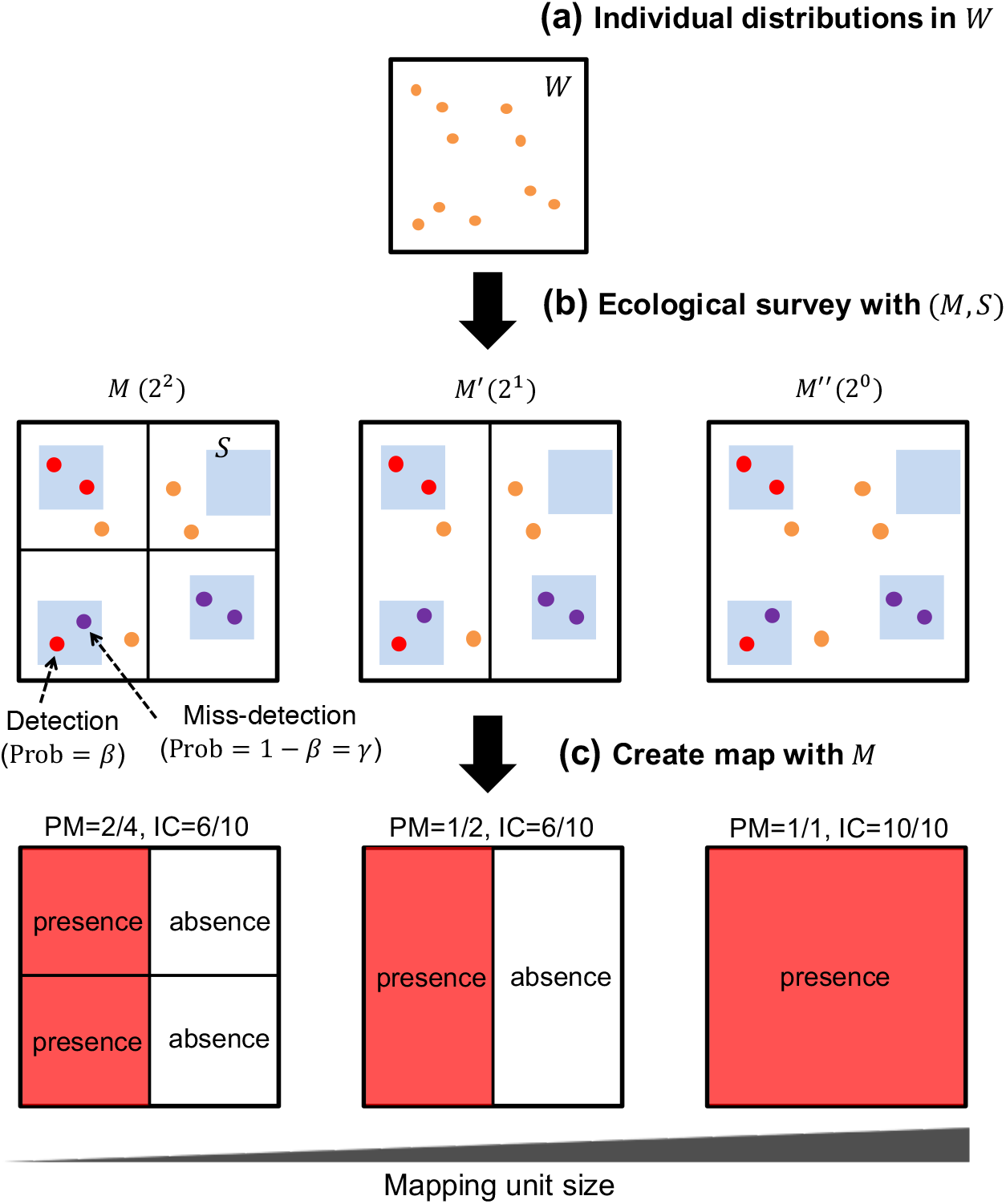
(color online) Ecological survey scheme within the observation window *W*. (a) Given the individual distributions in the observation window *W*, (b) the ecological survey is conducted with a certain mapping resolution *M* (left scheme in the middle row, for example) and sampling unit *S = aM* (blue regions). Each column represents the result of an ecological survey with a different mapping resolution *M (M′, M″)*. 2^2^ (2^1^, 2^0^) in the parentheses represents the number of mapping units within the observation window, (c) A presence/absence map is created using the survey outcomes. If at least one individual is found in a mapping unit (middle row: represented by red point), regardless of whether other individuals situated therein were detected or missed (represented by purple or orange points), the unit is mapped as presence, absence otherwise. In this step, the PM fraction and IC fraction (see main text for the definitions) are calculated by simply counting the number of presence patches or the number of individuals situated within the (mapped) presence patches. Although the same individual distributions and survey outcome are used, the map output changes if different mapping resolutions are used.

#### 2.2.2 Modeling the ecological survey

Here, we model the ecological survey with the three main assumptions listed above. Let, on average, *N* individuals of a species be distributed over a given window *W*, which is the region under concern (i.e., *N* = *N(W)*). The manner of individual distribution follows either the homogeneous Poisson process or the Thomas process. The resolution of the presence/absence map is defined by the scale of mapping unit *M*, and every mapping unit is sampled with the sampling unit *S*(Fig. 3). The survey is associated by the sampling error for each individual at a probability *γ*:= 1 *— β*, which is the probability at which individuals are not detected despite being present, and where, *β* is the detectability of an individual. For simplicity, we assume that the area of each mapping unit is 1, 2, 4,*…*, or 2^ra^ times smaller than the area of a given window *W*. Let *v(X*) be the area of a region *X*. With the definitions detailed above, we obtain

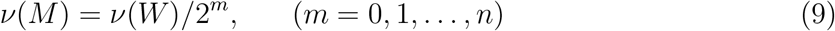

where, the superscript *m* represents the number of subdivisions of the window *W*. From Eq. (9), the number of mapping units within a given window *W*, is

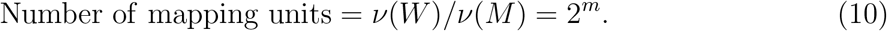

As the record for each mapping unit is based on an survey within the mapping unit, we obtain

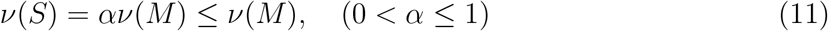

where, *α* is the sampling density within a mapping unit. Combining Eq. (9) and Eq. (11), we obtain

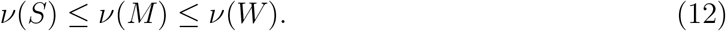

Let the intensity of the points within a given window *W* be [22,25]

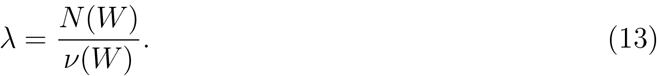

As we noted above, the parameters for the Thomas process are chosen so as to satisfy Eq. (8).

### 2.3 Assessing accuracy of presence/absence map

Given the spatial point pattern, sampling density, *α*, detectability of an individual, *β*, and scale of mapping unit, *M*, we calculate two main quantities of the ecological survey. That is, the presence mapped fraction (PM fraction), and, the fraction individuals covered by presence mapped patches (IC fraction):

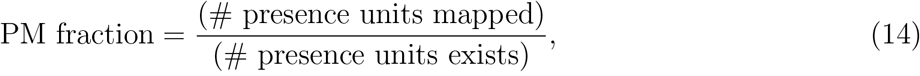

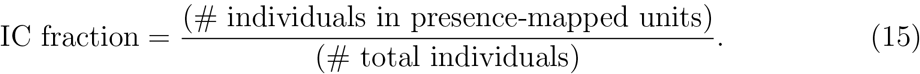

The PM fraction is the fraction needed to map presence units correctly and it relates to the probability of false negatives (i.e., 1 *—* (PM fraction) is the probability of the false negative in the map output). The IC fraction connects the PM fraction with population abundance in the observation window *W*, although this measure has not been used previously to our knowledge. For example, let us assume that we find the values of the PM and IC fraction are 0.8 and 0.95, respectively, given a survey scenario. In that situation, although we do not know where the individuals are located exactly, we expect that 95% of the total individuals in the observation window are situated within the presence mapped units. Therefore, the IC fraction also provides useful information for conservation. Examples of the PM and IC fraction values are given in Fig. 3. It is also expected that the difference between the average PM fraction and IC fraction increases as the degree of clustering in the distribution patterns increases, since the number of individuals in a mapping unit is biased to certain (moderate-sized) mapping units and such sites are more likely to be mapped as presence.

The false negative is often a concern in ecological monitoring to estimate how the monitoring is accurate (e.g., [13]). Nevertheless, we will apply the PM fraction to the following analysis to facilitate a comparison of the two measures, since the PM and IC fractions have similar forms as we will see below. As noted above, however, the degree of the false negative is directly evaluated from the PM fraction.

The presence mapped fraction is obtained by

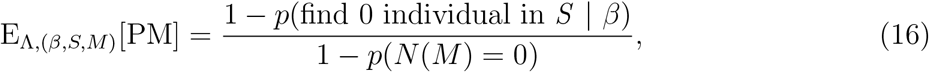

where, A indicates the underlying point pattern. On the other hand, the fraction of the total individuals situated within presence-mapped units is described as

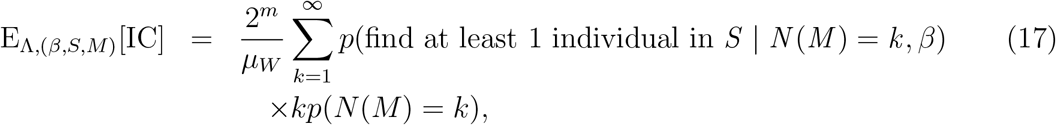

where, *2^m^* is the number of mapping units as Eq. (10). Since the IC fraction is rather cumbersome to derive analytically for the Thomas process, we only provide an analytical expression of the IC fraction for the homogeneous Poisson process, and give numerical results for the Thomas process.

### 2.4 Numerical settings

In addition to the IC fraction of the Thomas process, we conducted numerical simulations to confirm the validity of our analytical results using our own C code (available upon request). Implementing numerical simulations is straightforward by taking the first two steps (a) and (b) as shown in Fig. 3, and calculating the PM and IC fraction values by counting the detected habitats and individuals therein. We repeated this simulation 1000 times to obtain the 5, 25, 50, 75, and 95 percentile values. We set the observation window to 1024m *x* 1024m, and mapping unit was 2^1^, 2^2^, *• • •*, 2^17^ times smaller than the observation window. We set the sampling density and detectability to 0.5 and 0.9, respectively. The other parameter values were the same as in Fig. 1.

## 3 Results

### 3.1 Ecological survey with individual distributions based on the homogeneous Poisson process

Where individuals are distributed in space based on the homogeneous Poisson process, presence mapped fraction from Eq. (16) is

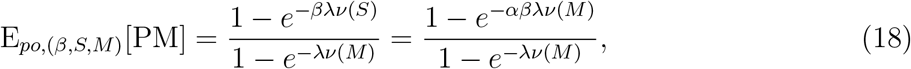

where, the equality *v (S) = αv (M)* is used. Eq. (18) has rather simple form and, thus, we can easily see the parameter dependence. The intensity of the points *λ* (Eq. 13) defines the average number of individuals existing within a given the observation window, *W*, and since 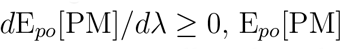 increases as the average number of individuals increase, and vice versa. Especially, when the intensity becomes ⋋ → ∞, E_po_[PM] becomes 1 regardless of the scale of mapping units. Intuitively, as the sampling density *α* and detectability *β* increase, E_po_[PM] increases, and vice versa. The asymptotic behavior *M* → 0 of Eq. (18) is obtained by expanding about *v(M)*

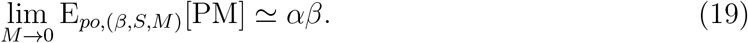

Since the zero probabilities *p(N(S*) = 0 | *β*) and *p(N(M*) = 0) approach to 0 as *M → W* given a sufficiently large observation window, we obtain

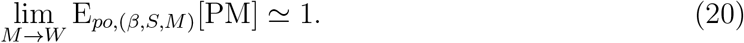

These results show good agreement with the numerical results (Fig. 4a).

**Figure 4:**
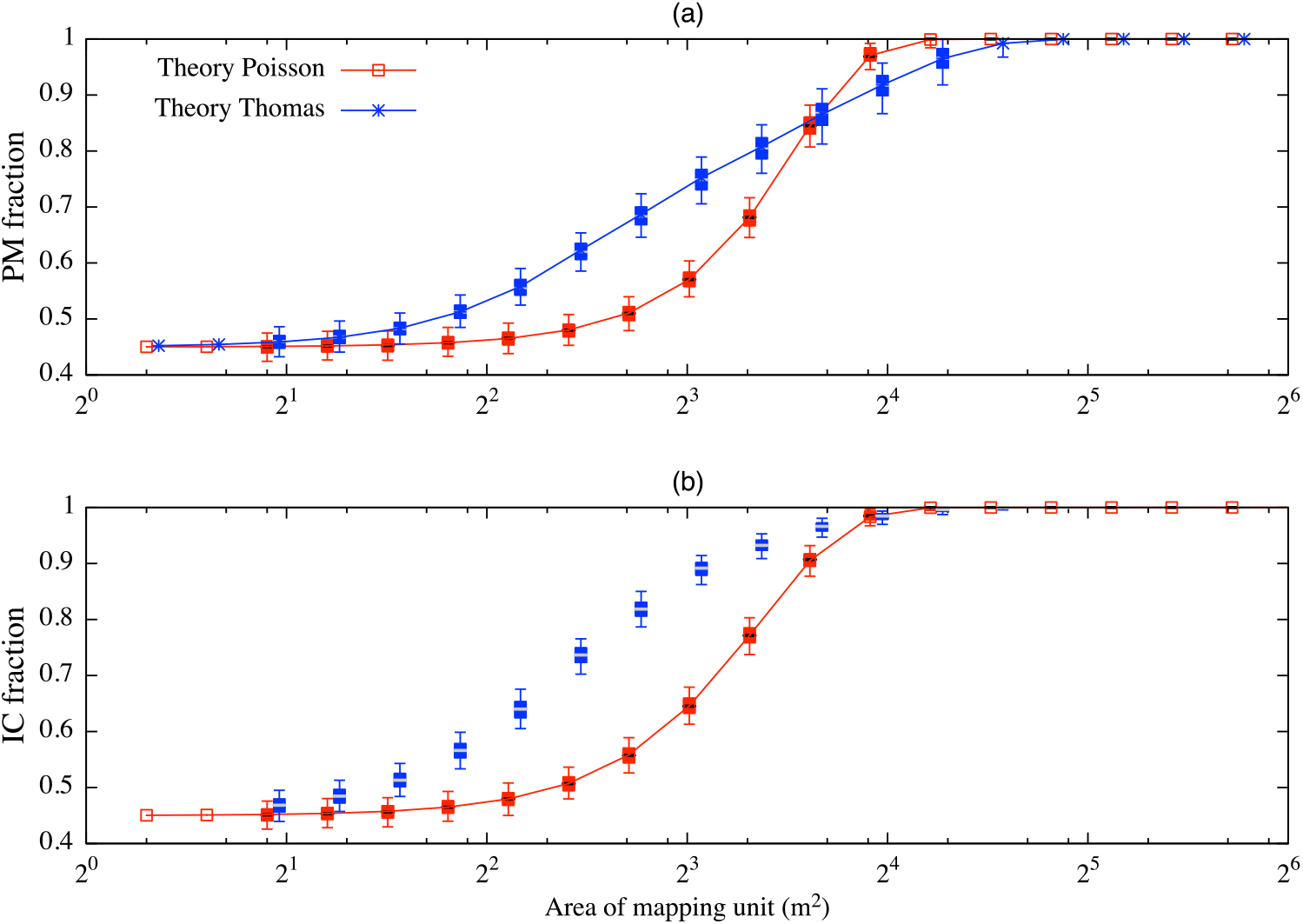
(color online) Analytical and simulated (candlestick) values of (a) the presence mapped fraction (PM fraction); and (b) the fraction of individuals covered within presence mapped patches (IC fraction) across mapping unit scales. *x*-axis is the area of mapping unit (m^2^). Each candlestick shows, from the bottom, 5, 25, 50, 75, and 95 percentile values of 1000 simulation trials. The values of the sampling density and detectability are α = 0:5 and β = 0:9, respectively. The other parameter values are the same as in Fig. 1.

For the homogeneous Poisson process, we can derive an analytical form of the average fraction of individuals covered within presence mapped patches (IC) as follows:

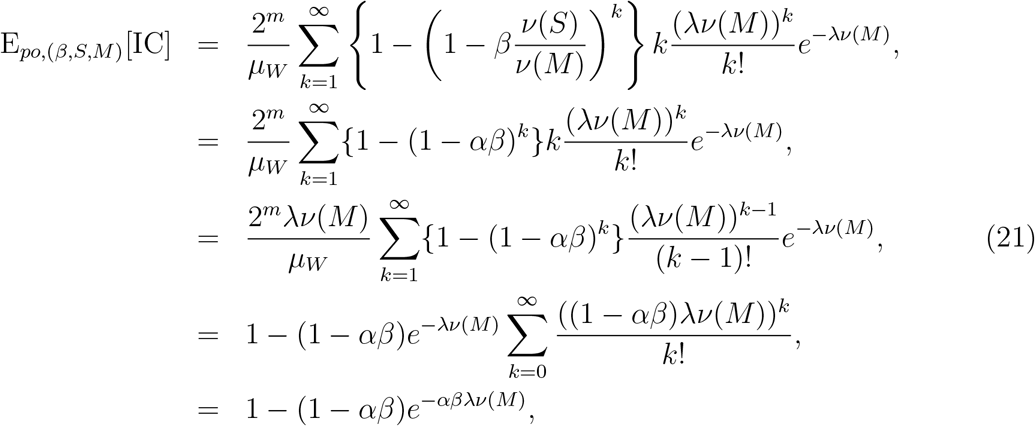

where, on the first line of right hand side, 2^m^ is the number of mapping units within the given window *W*, inside of the curly brackets is the probability that none of *k* individuals are detected by a survey given a mapping unit *M*, and the remaining term is the expected number of individuals within the mapping unit. The second line is obtained by using the fact *v(S*) = *αv(M*). To derive the fourth line, we used *μ_W_* = 2^m^*⋋v(M)*, and this equality is easily obtained by Fqs. (2) and (9). The dependences of the parameters ⋋, α, and *β* are qualitatively the same as those of Fq. (18). In addition, the asymptotic behaviors of Fq. (21) are equivalent to Fqs. (19) and (20). Fig. (4b) confirms the analytical evaluations of E_po_[IC].

Difference between the PM and IC fractions appears with an intermediate mapping unit size (see Fig. A.1 for a direct comparison), but the deviations are relatively small and these curves have similar forms. As mentioned above, it suggests that the degree of clustering is not large.

**Figure A.1:**
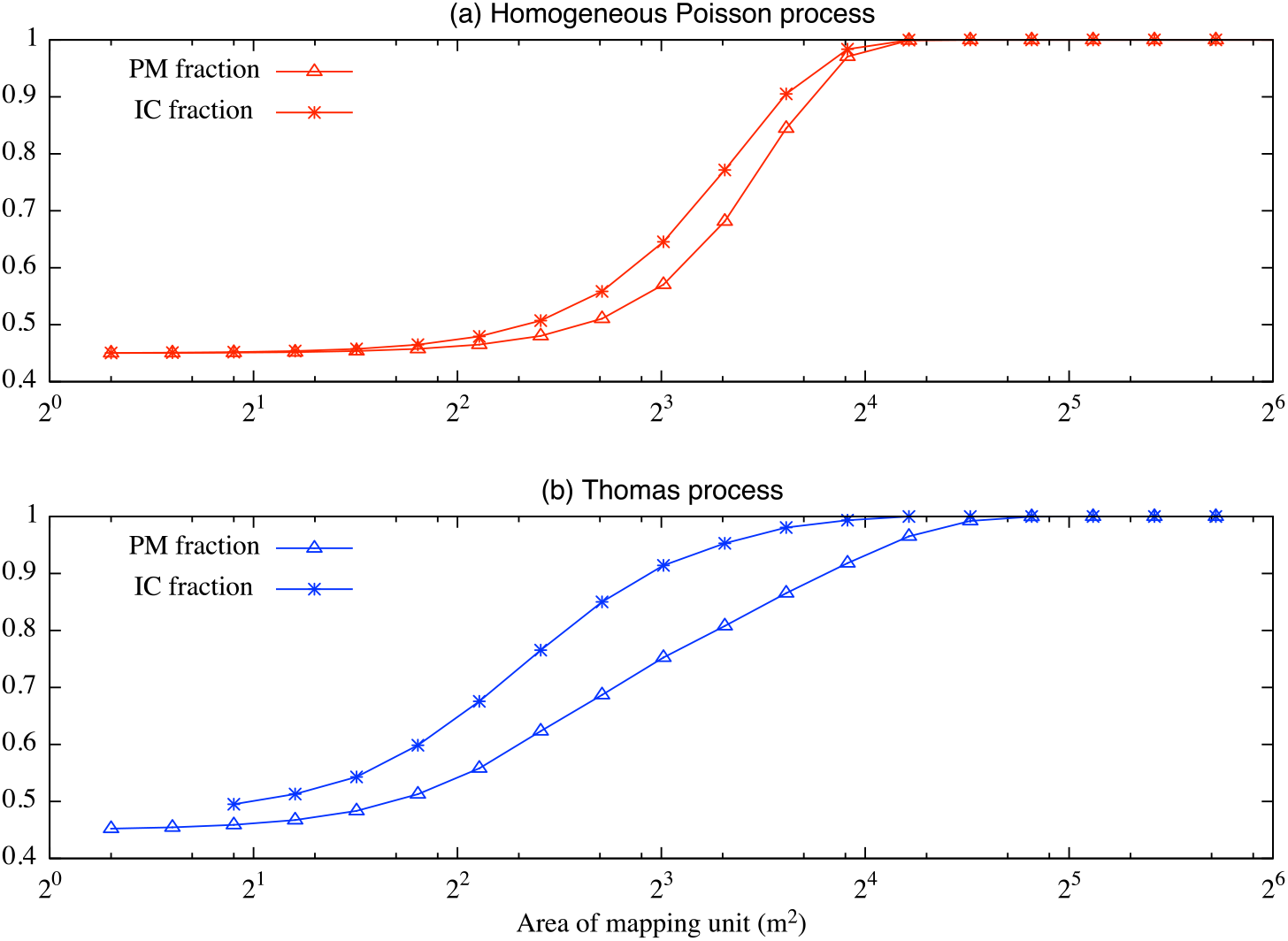
(color online) Analytical and simulated (only IC fraction of Thomas process) values of (a) the homogeneous Poisson process; and (b) Thomas process. All the values are the same as in Fig. 4 in the main text, but different presentation to facilitate the comparison of PM and IC fractions of each process.

### 3.2 Ecological survey with individual distributions based on the Thomas process

Here we consider the situation where individuals are distributed according to the Thomas process. By Fq. (16), we calculate the presence mapped fraction for the Thomas process:

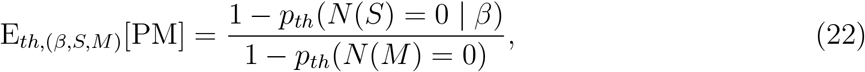

where, the probability of each event of the Thomas process is obtained by the pgfl Fq. (4): 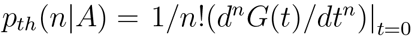. Therefore, *p_th_(N(A) = 0)* is

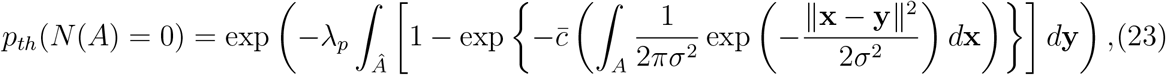

where Â is the surrounding region of *A* where parents potentially provide daughters to the region A. Specifically, the second term inside the square brackets for *p_th_(N(M*) = 0) in Fq. (22) becomes exp 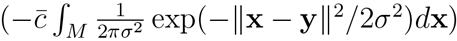 and that of 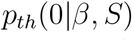 becomes exp 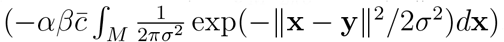, due to the sampling density and the detectabil-ity. Although Fq. (22) with Fq. (23) is not easy to interpret, we can calculate its asymptotic behavior in a similar manner to the derivations of Fqs (19) and (20):

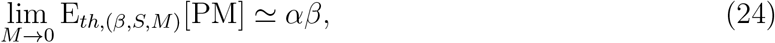

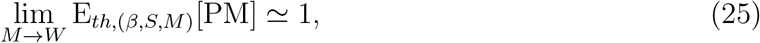

They are equivalent to the asymptotic behaviors of the homogeneous Poisson process Fqs. (19) and (20). Fig. (4a) plots analytical and numerical results, showing the theoretical value has a good agreement with the numerical calculation.

To obtain an explicit form for IC fraction of the Thomas process is cumbersome as the pgfl of the Thomas process Eq. (6) is rather complex. Therefore, we only show the numerical value for the IC fraction of the Thomas process (Fig. 4b). The IC for the Thomas process increases faster than Fq. (18) as the mapping scale increases. The asymptotic behavior shows similar trends to the other results.

As in the case of the homogeneous Poisson process, difference between the PM and IC fractions appears between the ranges where the asymptotic behavior occurs (Fig. A.1). However, the deviations are larger for the Thomas process, and it occurs with a wider range of the mapping unit size compared to the results of the homogeneous Poisson process. This is an effect of clustering distributions as discussed above.

## 4 Discussion

By explicitly accounting for the spatial distribution patterns of individuals through spatial point processes (SPPs) and multiple spatial scales of field survey, we develop a theory for ecological survey to map individual distributions. The theory quantifies two metrics, the presence mapped fraction (PM fraction) and the fraction of individuals covered by the presence mapped patches (IC fraction), and thus allows us to predict the outcome of an ecological survey under certain survey designs. When both the sampling density *α* and the detectability *β* are not equal to 1, we find a tradeoff between the value of the PM and IC fractions and the resolution of the map. The PM and IC fractions show the equivalent asymptotic behaviors for both the homogeneous Poisson process and the Thomas process where *αβ* and 1 are the outcomes of the small and large asymptotic limit of mapping units, *M*, respectively. In fact, these asymptotic limits occur generally for any distribution patterns if an observation window holds a sufficiently large number of individuals, which ensures that the probability of missing all individuals approaches zero. The limit of all these asymptotic behaviors are understood as follows: as the mapping unit scale becomes sufficiently small, each mapping unit can hold at most one individual. In such a situation, the probability of detecting the single individual is *αβ*. The asymptotic behavior suggests that there is a certain scale of the mapping unit above or below which the performance of an ecological survey does not change. Thus, in practice, we need to choose a scale of mapping units between these limits.

The PM fraction of the Thomas process first increases faster than that of the homogeneous Poisson process, because the Thomas process produces mapping units holding clustered individuals which are more likely to be found than mapping units holding a smaller number individuals. On the other hand, the PM fraction of the homogeneous Poisson process approaches to its asymptotic limit faster than that of the Thomas process. This is due to the fact that the Thomas process also produces mapping units holding a few individuals which has a smaller chance to be mapped as presence until the mapping unit becomes sufficiently large. This explanation may be used for any distribution patterns. For example, if individual distributions show highly clustered patterns, the PM fraction quickly increases at first and becomes gentle as the PM fraction approaches to the asymptotic value 1.

Spatial extension of the ecosystem that SPPs accounting individual aggregations describes could be large enough to cover a wide range of spatial scales. For example, Azaele et al. [24] showed that a Thomas model fitted to the distribution map of British rare vascular plant species (see the detailed description of the data set [29]) with three coarse resolutions (40000, 10000, and 2500 km^2^) can outperform many existing spatially-implicit models in terms of the down-scaling predictions of the species occupancy probability. In addition, Grilli et al. [30] showed that a special case of the Poisson clustering processes, a group of the point processes where parents locations are followed by a Poisson process [25] such as the Neyman-Scott process, recovers the species-area relationship at a local scale to continental scale as predicted by various existing models (e.g., [31]). Hence, even though we used a observation window *v(W)* = 1024m x 1024m as an example, it can be generalized by changing its scale and the sampling intensity. In addition, it is worth noting that albeit individuals of most species are typically aggregated [32, 33] the Thomas process could be approximated by the homogeneous Poisson process under a certain condition: when the intensity of individuals is large, the PM fraction of the Thomas process comes close to that of the homogeneous Poisson process 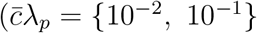 in Fig. A.2). This is due to increased parent intensity decreasing spatial heterogeneity over the region concerned, suggesting potential applicability of the simpler model to an abundant ecosystem.

**Figure A.2:**
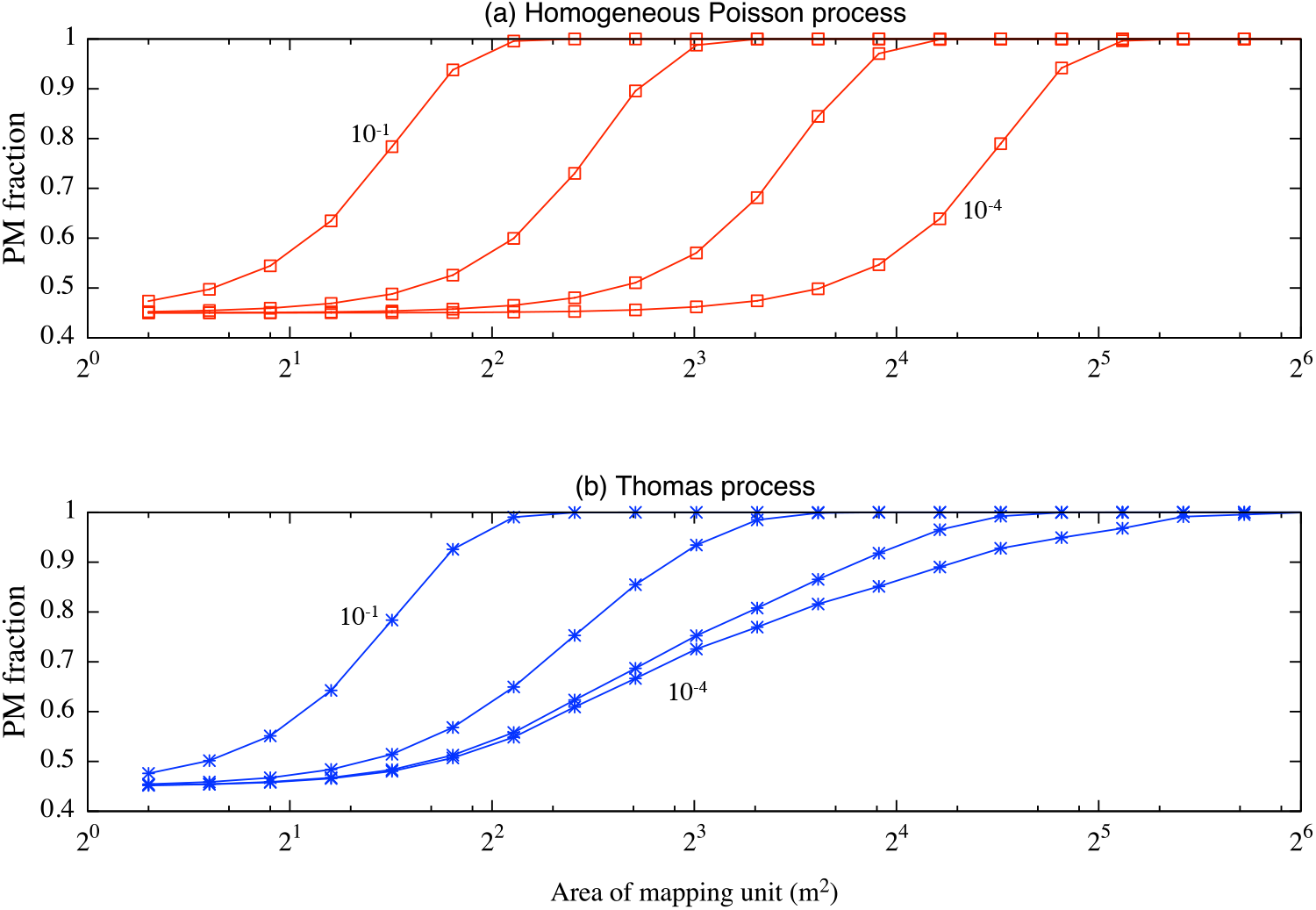
Effect of the intensity 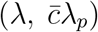 in the observation window, *W*, on the theoretical presence mapped (PM) fraction, Eqs. (18), (22). The intensity of the Thomas process is manipulated by changing the parent intensity *λ_p_*. Individual distribution patters are according to the (a) Homogeneous Poisson process and (b) Thomas process. For the Thomas process, the curves for PM fraction converge as the intensity becomes small, and come close to the corresponding curve of the homogeneous Poisson process as the intensity of the Thomas process increases. This is an efect that the increased parents intensity decreases spatial heterogeneity over the concerned region. For both panels, the order of the intensity monotonically decreases from left to right.

For simplicity, we consider a situation where each mapping unit is sampled with the same sampling density, *α*, and detectability, *β*, and the location of the sampled unit within a mapping unit is chosen randomly. These are rather idealized assumptions and may be further generalized. For example, it may be reasonable to assume that the sampling density, *α*, and the detectability, *β*, become almost 1 at a certain fine scale of the mapping unit. Although such a fine scale may not be achieved because of budgetary constraints, explicitly taking into account the spatial effect on *α* and *β* gives us better understanding about the fine scale of asymptotic behavior. In practice, the location of the sampling unit may be determined by more strategic manner depending on ones purpose. Indeed, previous studies had proposed several sampling strategies which emphasize, for example, a spatially contiguous placement of the sampling units to correctly capture ecological patterns (e.g. [34]), a systematic placement to efficiently reflect spatially structured ecological processes [35, 36], or a representative design for major environmental gradients to maximize per effort information of organism’s distribution [37, 38]. While these strategies have been compared empirically using actual dataset (e.g. [36]), the developed theory in this paper may provide a theoretical base to evaluate the effectiveness and efficiency of such purpose-dependent sampling strategies.

### Connection to occupancy area and population abundance

Presence/absence maps are often used to estimate the occupancy area or population abundance [24, 39, 40]. Since our map contains estimated inaccuracy, we need to consider this effect to estimate these quantities. In our framework, occupancy area is straightforward to obtain using the number of occupancy units. The number of occupancy units is calculated from the PM fraction and the presence/absence map from an ecological survey, since we know the relationship: (# presence mapped units) = F[PM] *x* (# total occupancy units). We can also derive the number of occupancy units using the following relationship: (# total occupancy units) = 2*^m^*(1 – *p*(*N(M)* = 0)), where 2*^m^* is the number of mapping units. Unlike the tradeoffs between mapping resolution and the PM or IC fraction, this estimation is improved by a survey with finer-scale mapping units since the shape of an occupancy region is better mapped by a finer resolution. However, this effect has a finite limit.

Population abundance is also estimated using the fact that each mapping unit at most can hold one individual at a sufficiently small mapping scale. At this limit, the estimated number of the total occupancy units corresponds to the total population *N(W)*. In fact, we see the following relationship, for example with the homogeneous Poisson process, *N(W)* = lim_M→0_2^m^(1 – *p*(*N(M)* = 0)) = *λv (W)*, by the equality 2^m^ = *v(W)/v(M)* and the same expansion as in Eq. (19). *λv(W)* is the unbiased estimator of the total population due to the definition of Eq. (13). This estimation, however, requires rather small mapping unit sizes and, therefore, it is seldom practical. Therefore, a limited number of sampling trials to estimate the probability *p*(*N(M)* = 0) may be required.

### Application to conservation/ecosystem management

For decision making regarding field survey designs, mapping resolution must be determined to balance the accuracy (i.e., the PM and/or IC fraction) and resolution of the map. Our results show that map accuracy is improved with larger mapping resolution. However, presence/absence maps with too coarse a resolution are not useful for many ecological studies and conservation/management practices. In addition, Takashina and Baskett [7] showed that fisheries management using a coarse management unit inevitably increases inefficiency. Therefore, it may be reasonable to start by first determining the required map accuracy, and second by finding the finest possible mapping resolution at which an expected PM or IC fraction satisfies the requirement. As an example, let us discuss a simple and ideal situation where we have estimations of each parameter value and the population abundance within an observation window. For simplicity, we assume that all the parameter values are the same as in Fig. 4, and the target species has a clustering distribution pattern, which is described by the Thomas process (corresponding to the situation shown in Fig. 4b). Let us further assume that the situation where we found 55% of the population in the observation window must be protected to satisfy a 95% chance of persistence over the next 100 years through population viability analysis. Therefore, the minimum requirement of the ecological survey is to produce a presence/absence map with at least 55% of the total population covered within the presence mapped units. By making use of Fig. 4b, we then find that the size of the mapping resolution *M* needs to be about 64m^2^ or larger to meet this requirement, of which an expected value of the IC fraction is F_th_[IC] = 0.57. That is to say, we are expected to get a presence/absence map in which 57% of the total population is situated in the presence mapped units. If the implied mapping resolution is too coarse to apply, then managers must improve either the detectability, *β*, or sampling density, *α*, to meet the required IC fraction and produce a map with a finer resolution.

Of course this example oversimplifies the ecological survey program, since we often do not have parameter values of a target species. However, the concept discussed above is rather general and hence applicable to wide variety of ecological surveys. The core of this idea is to clearly set the feasible goal, with time and budgetary constraints, of conservation practice or ecological study in advance.

In practice, the developed theory should be, to an extent, complemented by an estimation of the existing number of individuals within the observation window, *W*, in advance, since the intensity affects PM and IC fractions (Fig. A.2). An estimation of the population abundance could be done by using historical record, demographic data or surrogate data depending on availability of information. Statistical and theoretical methods such as species distribution modeling [41] estimating the occurrence of plant species across scale [24, 42] or predicting the population abundance in a coral reef environment [43] may complement these methods. Conducting a pilot survey is one alternative way to estimate the population abundance with a required estimation accuracy. Takashina *et al*. [28] recently developed a framework for the pilot sampling providing a required minimum sampling effort to satisfy the required accuracy. Complemented by these steps, the theory developed here has a potential to significantly improve survey frameworks.

## Acknowledgements

We thank M. Akasaka, B. Stewart-Koster, T. Fung, R.A. Chisholm, L.R. Carrasco and S. Azaele for their thoughtful comments and discussions. NT and BK were funded by the Program for Advancing Strategic International Networks to Accelerate the Circulation of Talented Researchers of the Japan Society for the Promotion of Science. They acknowledge the support for coordinating the research program from Dr Yasuhiro Kubota and Dr James D. Reimer. BK was also funded by The University of the Ryukyus President’s Research Award for Leading Scientists.

